# Self-similar tip growth links exocytosis profile with cell wall shape

**DOI:** 10.1101/2022.04.26.489612

**Authors:** Kamryn Spinelli, Chaozhen Wei, Luis Vidali, Min Wu

## Abstract

Exocytosis plays a crucial role in regulating the growth and migration of filamentous tip-growing cells. We present a mathematical framework that infers the spatial profile of exocytosis from the cell morphology in self-similar growing cells that elongate while preserving their apical domain shapes. By applying the framework to cell wall outline data from experiments across walled cell species, we find that while tapered cells have their exocytosis concentrated at the apex, cells with flatter tip shape beyond a threshold require exocytosis to peak in an annulus region away from the apex.

---

In plants and fungi, the cell wall boundary defines the cell morphology and sustains the internal turgor pressure resulting from osmosis [1]. In tip-growing filamentous cells, such as in root hairs [2], moss protonemata [3], pollen tubes [4], and fungal hyphae [5], which are one cell wide, cell walls need to extend along one direction to optimize the migration speed. How this growth mode is regulated is of critical importance for plant and fungus development and survival.

Without perturbing external environmental cues, the filamentous cell wall in its apical domain is approximately axisymmetric and extends along its axis of symmetry [3, 5, 6]. Many filamentous cells can elongate while preserving their apical-domain shape, called self-similar tip growth [7]. This specific morphogenetic process results from the spatial patterning of two factors: secretion of new wall material, cellulose synthase protein, and wall loosening factors via exocytosis; and mechanical cues in the cell wall surface induced by the turgor pressure [1].

Previous work on filamentous growth has developed mathematical models to partially explain how mechanical cues and secretion patterns regulate the cell shape during wall expansion by approximating the cell wall as a thin-shell surface [7–14]. These theoretical models can be classified into visco-plastic [8], viscous fluid [9–12], and growth-elastic models [7, 13, 14]. While the local wall-surface growth rate is assumed to only respond to wall tensions in the visco-plastic model, it is determined by the secretion pattern in the viscous fluid model. However, in the fluid models, the cell-wall viscosity must increase unboundedly in regions far away from the tip end to generate realistic cell profiles. The growth-elastic models explain cell wall expansion as the combined effect between irreversible growth and elastic deformation. However, they have not explicitly connected the secretion pattern with the surface growth process.

The study of secretion patterning in tip-growing cells is crucial because it connects the internal cellular processes such as vesicle transport and exocytosis to cell morphogenesis [15]. In particular, limited by current experimental approaches [16], a consensus has not been reached on the spatial distribution of exocytosis in pollen tube growth [17]. While exocytosis is generally thought to concentrate at the tip end, measurements of vesicles in the vicinity of the apical cell membranes suggest that the exocytosis is patterned at an annulus region centering on the tip [18, 19].

For more than a century [20–22], we have been able to track the relative cell-wall surface expansion rates 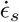 and 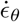 along the meridional and circumferential direction, respectively, through measuring the rate of tangential velocity *v*_*s*_ = *∂s/∂t* and local cell width *r* via [6, 22] (see Fig.1a)

**FIG. 1:**
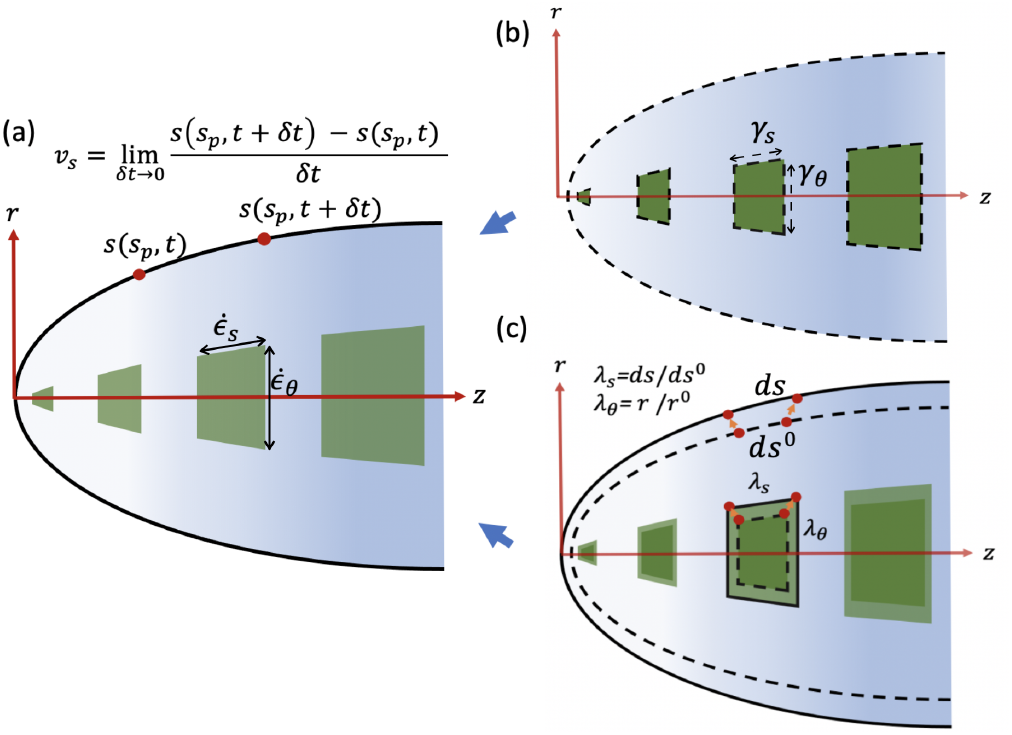
The observed wall expansion (a) is a combined effect of wall expansion of the relaxed, unturgid, configuration (b) and an elastic deformation (c).

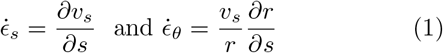

where *s* is the meridional distance from the tip end. In root hair growth, the experimental tracking of wall surface expansion [6, 22] shows that the cell wall expands with highest rates 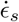 and 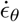 in an annulus region around the tip end. However, these measurements cannot be used directly as the readout of the spatial profile of exocytosis because the wall mechanics also influences the wall surface growth [1]. Rather, it is commonly assumed that the exocytosis profile is consistent with vesicle accumulation underneath the apical domain [23]. Nevertheless, neither experimental nor theoretical studies have reported whether the exocytosis itself indeed distributes at the very apex or annularly around the tip end in root hairs.

This letter presents a mathematical model that describes the cell wall surface growth as a combined effect from the elastic deformation and the irreversible expansion in response to the local exocytosis and elastic strain. The distinguishing feature of our model is the explicit reliance on the cell’s relaxed configuration without turgor pressure (we define “relaxed” by “unturgid” henceforth) and the connection between the unturgid and turgid configurations by an elastic deformation. By exploiting the self-similarity condition ([24], see below), we formulate a well-defined initial value problem to solve the exocytosis profile from the cell shape data and apply the framework to multiple cell species.

Treating the cell wall as a thin shell of revolution generated by its meridian, we assume that at any time point *t*, one material point *s*_*p*_ along the meridian is mapped to two configurations, one turgid state (*s*(*s*_*p*_, *t*), *r*(*s*_*p*_, *t*)) and one unturgid state (*s*^0^(*s*_*p*_, *t*), *r*^0^(*s*_*p*_, *t*)) where *s* and *s*^0^ are the meridional distances from the tip end and *r* and *r*^0^ are the local cell widths in the turgid and unturgid states, respectively. We describe the growth of the unturgid surface path by

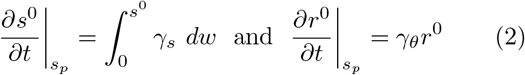

where *γ*_*s*_ and *γ*_*θ*_ are the expansion rates of the unturgid cell wall surface in the meridional and circumferential directions respectively (Fig. 1b). The first part of the equation comes from the fact that the rate at which a point of wall material at arclength coordinate *s*^0^ moves away from the tip end is given by summing the local meridional expansions from the tip end to that point. The second part of the equation comes from the fact that the circumference of the circle traced by rotating the material point *s*_*p*_ around the axis of revolution is *ℓ*_*r*_ = 2*πr*^0^. It follows that *∂ℓ*_*r*_*/∂t* = 2*π*(*∂r*^0^*/∂t*) = *γ*_*θ*_*ℓ*_*r*_ = 2*π*(*γ*_*θ*_*r*^0^).

The unturgid and turgid configurations are connected by the elastic stretches (Fig. 1c)

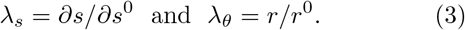

That said, we describe the cell wall surface area growth coming from two contributions: 1) expansion rates of the unturgid surface patch and 2) the stretch ratio due to elastic deformations *λ*_*s*_ and *λ*_*θ*_. In this regime, we assume that the local expansion rates of the unturgid surface patch are induced by the exocytosis rate *γ* and the elastic strains along the meridional and circumferential directions:

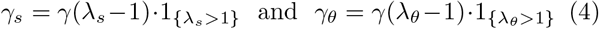

where 1_*χ*_ is the characteristic function of the set *χ*.

In self-similar tip growth (the apical-domain shape is in a steady state), the local expansion rates of the material points need to satisfy a constraint 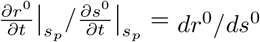 where *r*^0^(*s*^0^) describes the steady-state outline of the unturgid configuration. We name this constraint the self-similarity condition, which has been derived previously in [24]. Combining this condition with Eqs.(2) and (4), we derive

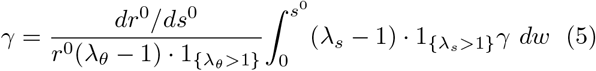

which enables us to solve *γ* given the steady-state cell geometry *r*^0^(*s*^0^) and elastic stretches *λ*_*s*_ and *λ*_*θ*_.

Henceforth we consider the apical domain from the tip end *s*^0^ = 0 to a limit point *s*^0^ = *s*^*b*^, which we call the shank boundary, along the meridian in which the conditions *λ*_*s*_ *>* 1, *λ*_*θ*_ *>* 1 and *dr*^0^*/ds*^0^ *>* 0 hold. Under these conditions, one can obtain *γ*^*′*^ (*s*^0^) = *f* (*s*^0^)*γ*(*s*^0^) by differentiating Eq.(5) with respect to *s*^0^. With the condition 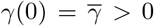 which sets the scale of the exocytosis profile, one can show the distribution *γ* is uniquely determined. See *f* (*s*^0^) and the detailed analysis in the Supplemental Material [25]. Importantly, one can show that 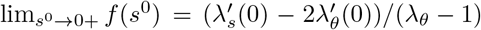). This shows that there is a local maximum of *γ* at the tip location *s*^0^ = 0 only when the elastic stretches satisfy the relation 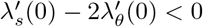, and vice versa.

We estimate the elastic stretch distributions by developing a thin-shell mechanics model. To describe the the cell wall mechanics, both wall-surface tension and bending moment may be considered [26, 27]. For simplicity, we suppose that the force 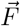 on the cell wall is dominated by the wall-surface tensions *σ*_*s*_ and *σ*_*θ*_ along meridional and circumferential direction respectively, and the turgor pressure *P* :

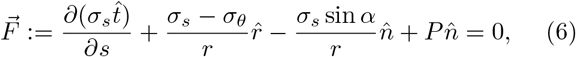

where 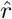 is the unit vector along the cell width, 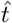 is the local unit tangent, 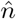 is the local unit normal vector towards the cell exterior, and *α* is the angle from 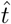 to 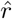. This vector equation can be derived from 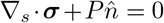 where 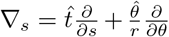 and 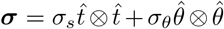, *θ* being the azimuth with respect to the axis of symmetry. The widely used Young-Laplace Law [7–14]: *κ*_*s*_*σ*_*s*_ + *κ*_*θ*_*σ*_*θ*_ = *P* and *κ*_*θ*_*σ*_*s*_ = *P/*2 (where *κ*_*s*_ and *κ*_*θ*_ are the curvatures along the meridional and circumferential directions, respectively) can be derived from this equation.

To relate the wall-surface tension with the elastic stretches, we first consider the nonlinear elastic theories [7]. In particular, we consider the constitutive law

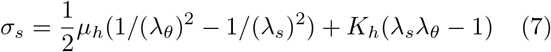

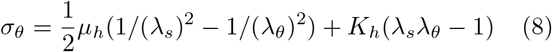

where *K*_*h*_ = *h × K* and *µ*_*h*_ = *h × µ* are the rescaled bulk modulus and shear modulus, respectively, *h* being the cell wall thickness in the thin-shell approximation. Later we will also consider the linear law 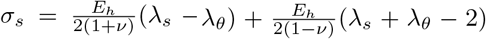 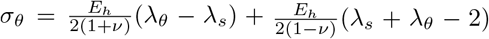 where *E* = *h E* and *ν* are the rescaled Young modulus and Poisson ratio respectively. These two laws are equivalent up to first order of small strains. The linear elastic law has been considered in [27].

At the tip end of the apical domain, we have the condition

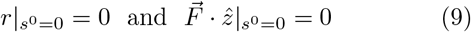

due to fixing the tip along *r*-direction and force-balance along the long axis *z*-direction 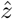. Similarly, at the shank boundary *s*^0^ = *s*^*b*^, we have

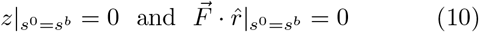

due to no displacement of the shank boundary along the *z −* direction and force-balance along the *r −* direction. Notice *z*(*s*) is constrained by *r*(*s*) by the geometric fact (*dz/ds*)^2^ + (*dr/ds*)^2^ = 1.

To summarize, the theoretical framework infers *γ* given the steady-state unturgid configuration *r*^0^(*s*^0^) on [0, *s*^*b*^] via Eq.(5). To simulate the elastic stretch *λ*_*s*_(*s*^0^) and *λ*_*θ*_(*s*^0^) distributions given *r*^0^(*s*^0^), we solve the turgid configuration *r*(*s*) and the mapping of arc-length coordinate *s*(*s*^0^) via Eqs.(3) and (6)-(10), where *λ*_*s*_ and *λ*_*θ*_ are computed from *s*(*s*^0^) and *r*(*s*) via Eq.(3). See Supplemental Material [25] for the solution procedure.

By rescaling *r*^0^ → *r*^0^*/L, r* → *r/L, s*^0^ → *s*^0^*/L, s* → *s/L, µ*_*h*_ → *µ*_*h*_*/*(*PL*), *K*_*h*_ → *K*_*h*_*/*(*PL*), *E*_*h*_ → *E*_*h*_*/*(*PL*) (where *L* = *r*^0^(*s*^*b*^) is half of the cell width at the shank boundary) and using the same notation for the nondimensionalized quantities, we end up having only two material parameters *µ*_*h*_ and *K*_*h*_ (or *E*_*h*_ and *ν*) in the model and the dimensionless cell shape outlines *r*^0^(*s*^0^) and *r*(*s*).

We first apply the framework to the moss *Physcomitrium patens* protonemata and *Medicago truncatula* root hairs that grow their tips approximately self-similarly. Since only the turgid cell outline data are available previously [6, 28], we first use the turgid cell outline *r*(*s*) to approximate the unturgid *r*^0^(*s*^0^). For each cell type (moss caulonemata, moss choloronemata, and root hair), we define its canonical shape *r*^0^(*s*^0^) by fitting multiple cell outline data using a “hyphoid” function *z* = (*πr/a*) cot(*πr*) [29]. We find that this hyphoid function fits the tip cell data rather well, with less fitting error than the ellipse function continued by a flat region [14]. See Supplemental Material [25] for the data-fitting details.

Applying our theoretical framework to each canonical cell shape *r*^0^(*s*^0^) with material parameters *K*_*h*_ = *µ*_*h*_ = 24 (we will later discuss the choice of *K*_*h*_ and *µ*_*h*_), we infer their exocytosis distribution *γ* (Fig.2). For moss protonema canonical shapes, we find the exocytosis distributes highest at the tip end (e.g., see Fig.2a for choloronema). In this case, the elastic stretches satisfy *λ*_*θ*_, *λ*_*s*_ *>* 1 and 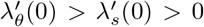 (Fig.2a). Combined with the analysis of the self-similarity condition Eq.(5), we have 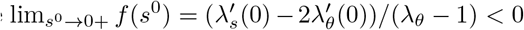, which means *γ*(0) at the tip end is a local maximum.

**FIG. 2:**
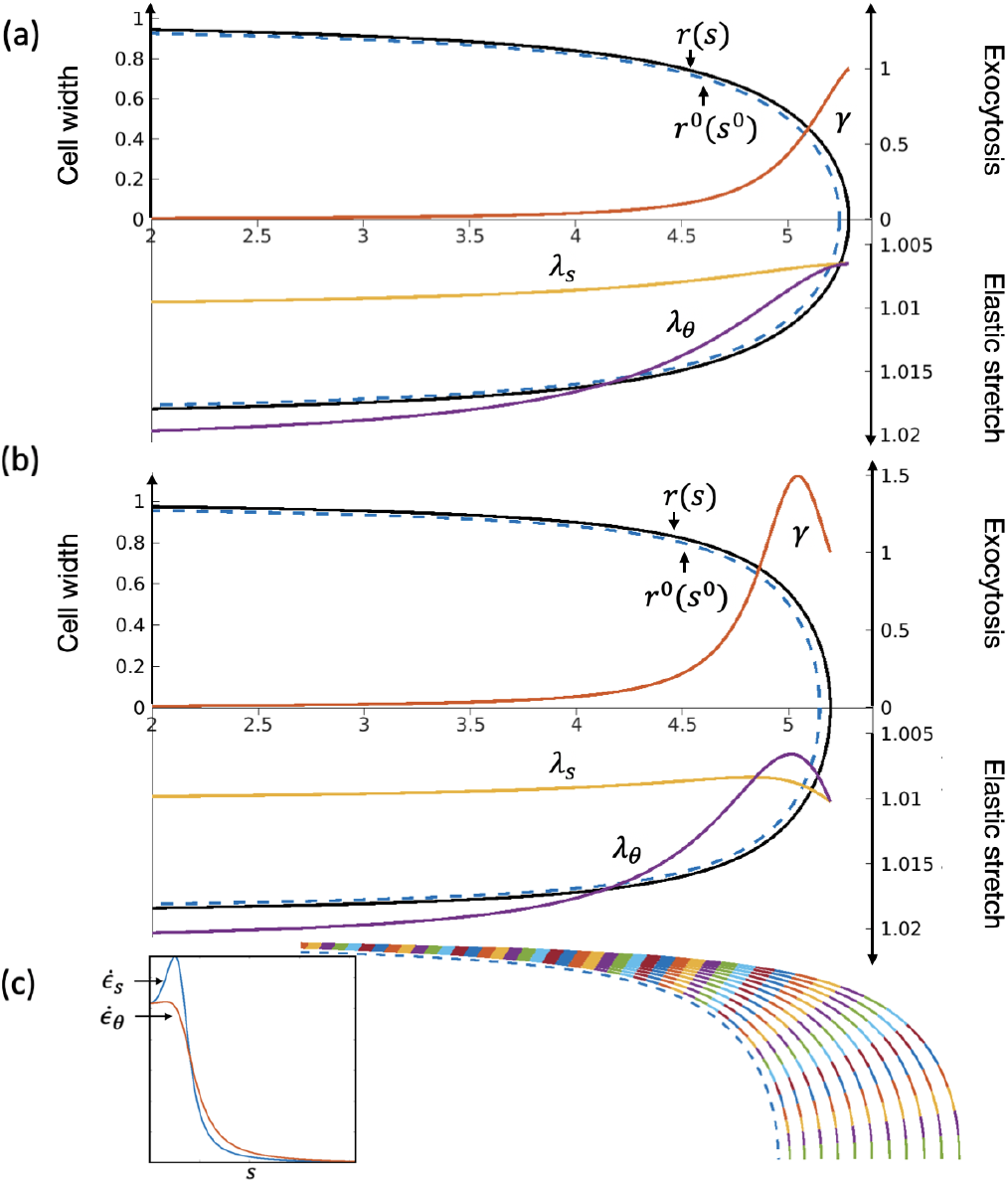
For the canonical shape of moss choloronema, the exocytosis profile *γ* is monotonically decreasing from the tip, with elastic stretches *λ*_*s*_ and *λ*_*θ*_ increasing from the tip (a). For the canonical shape of *Medicago truncatula* root hair, *γ* is non-monotone and has its global maximum away from the tip, with non-monotone *λ*_*s*_ and *λ*_*θ*_ (b). The predicted wall extension rates 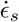 and 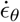 of *Medicago truncatula* root hair (inset) are computed from the simulation (c). They are also non-monotone, consistent with the measurements in [6].

In contrast, the root hair canonical shape has exocytosis peaking annularly around the tip end (Fig.2b). In this case, the elastic stretches satisfy *λ*_*θ*_, *λ*_*s*_ *>* 1 and 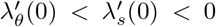. Thus lim_*s*_0 _*→*0+_ *f* (*s*^0^) *>* 0, which means *γ*(0) cannot be a local maximum. By assuming that lim_*s*_0 _*→∞*_ *γ* = 0 and *γ* is continuous, there exists a 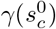 such that *γ*(*s*^0^) is the global maximum, as shown in Fig.2b. To compare with previous measurements on the cell wall extension rates 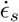 and 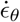 of *Medicago truncatula* root hair tips [6], we also simulate self-similar root hair tip growth by using Eqs.(2) and (4) to update (*s*^0^(*s*_*p*_, *t*), *r*^0^(*s*_*p*_, *t*)) using the inferred *γ* and solve the turgid configuration (*s*(*s*_*p*_, *t*), *r*(*s*_*p*_, *t*)) at each time step, and we compute extension rates 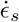 and 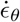 using Eq.(1) (Fig.2c). We find that 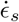 and 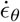 have their maxima away from the tip end (Fig.2c, inset), and this is consistent with the experimental measurements in Figure 4c-f in [6]. In addition, the location with the extension rate maxima does not present the highest elastic stretches. Altogether, our results indicate the measured non-monotonicity in extension rates of *Medicago truncatula* root hairs is due to the annular exocytosis distribution.

Further, we change the material parameters *K*_*h*_ and *µ*_*h*_ and the constitutive law of the cell wall elasticity. We find the above results do not change qualitatively under this change of elasticity theory and the mechanical parameters. See Supplemental Material [25] for details.

To link the exocytosis profile with the cell shapes, we obtain the transition of cell shapes from tapered to flat apical domains by changing the value of *a* in the hyphoid function *z* = (*πr/a*) cot(*πr*). As the apical domain becomes flatter, *γ* transitions from a monotone decaying function of the meridional distance to a profile that first increases and then decreases to zero. See Supplemental Material [25] for details and figures. The transitions of cell wall expansion rates 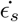 and 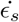 behave similarly. On the contrary, the elastic stretch distributions transition from monotonically increasing to a steady level to decreasing first, followed by an increase to a constant level.

We quantify the change of exocytosis profile versus the cell shape by measuring the length of the secretion window *w* (defined as the shortest arclength interval [*s*_1_, *s*_2_] containing half the total secretion) and the meridional position *l* of the highest exocytosis rate versus the radius of the tip curvature, *R* (see Fig.3). To eliminate the effect of the cell size, all the quantities are rescaled by the turgid cell width *r*(*s*^*b*^) at the shank boundary.

**FIG. 3:**
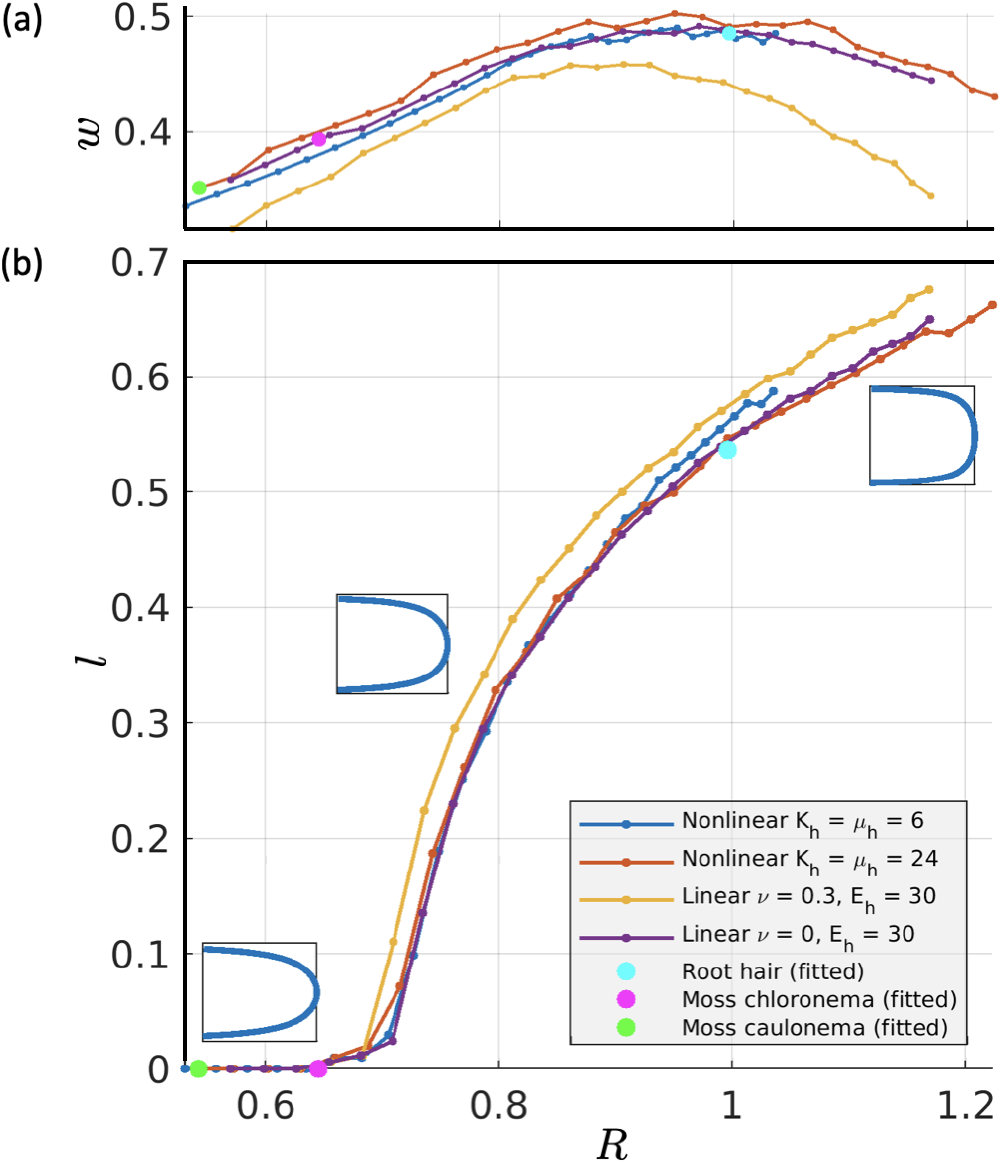
The width *w* of the exocytosis profile (a) and the meridional position *l* of secretion maximum (b) vs the radius of the tip curvature. The dependence of *w* and *l* on *R* is insensitive to the change of material parameters and the change of constitutive laws (see inset for the correspondence of each curve).

For relatively tapered cells, the tip radius of curvature *R* can be changed due to the narrowing or widening of the exocytosis profile concentrated at the tip. See Fig.3a for the range *R* ≤ 0.65 where the data points for moss chloronema and caulonema are located. Beyond a critical value *R* ∼0.65, the exocytosis peak has to move away from the tip to maintain flatter tip shape. See the takeoff of *l* when *R* ≥0.65 in Fig.3b. We have used linear and hyper-elastic models with different parameter values, but the quantitative relation from *R* to *l* and *w* is insensitive to the change of material assumptions (see Fig.3 and the caption).

Our results provide a geometric explanation for conflicting results in the location of fastest exocytosis, attributing them to differences in the cell shape [30]. For cell shapes that can be approximated by a hyphoid function, we find two strategies to reach a specific cell shape: adjusting the polarity of the secretion profile and moving the exocytosis “hotspot”. For tapered shapes close to moss protonemata (*R <* 0.65), perturbation of the secretion window width can change the tip shape. For flatter-tip cell shapes with *R >* 0.65, the secretion “hotspot” has to move away the tip end to generate a specific shape. Interestingly, the *Medicago truncatula* root hair canonical shape (*R ≈* 1) is associated with the widest secretion window (Fig.3a), suggesting that this shape is easier to maintain than slightly more tapered or flatter hyphoidfunction shapes. Given that more chemical energy is required to increase secretion polarity, the shape of *Medicago truncatula* root hair may be evolutionarily favored. To become even flatter (*R >* 1), a more focused exocytosis (smaller *w*) is required to maintain the shape (see the flat shape in the inset) and the shape change becomes more sensitive to the change of secretion “hotspot” (| *dR/dl* | becomes larger). This result provides a reason why no extreme flat filaments have been observed among different walled cell species.

Although an ellipse function continued by a flat region has been used to approximate the shape of the lily pollen tube [14], we found that it does not fit the root hair tips and moss protonemata shape well. We also found theoretically, that by changing the aspect ratio of the ellipse apical domain within the realistic range *R* ≤ 1, the highest secretion rate stays at the tip end (*l* = 0) and the change of tip curvature is highly sensitive to the change of secretion window width (| *dR/dw*| »1). This suggests that ellipse shapes are hard to maintain and may provide a clue why the lily pollen tubes undergo oscillatory shape evolution [11].

It is still unclear why multiple walled cell species are close to the outline *z* = (*πr/a*) cot(*πr*) in their averaged canonical shapes (e.g., fungal hypha [29], root hair tips, and moss protonemata). Our framework shows that the cell wall geometry uniquely determines the secretion profile. Still, the converse is not necessarily true, that is, whether a particular distribution of secretion uniquely determines a specific cell shape. From analyzing the self-similarity condition Eq.(5), the direct dependence on the shape function *r*^0^(*s*^0^) and its derivatives vanish at the tip *s*^0^ = 0, which makes it non-trivial to formulate a welldefined initial value problem for solving *r*^0^(*s*^0^) based on this steady-state condition. Future work will consider time-dependent shape evolution which may explain why the canonical shapes of multiple walled cell species distribute along the hyphoid function.

We thank Dr. Jacques Dumais for providing the root hair cell outline data from [6]. M.W. and K.S. acknowledge partial funding from the National Science Foundation- Division of Mathematical Sciences (NSF-DMS) grant DMS-2012330. L.V. acknowledges partial funding from the Division of Molecular and Cellular Biosciences (NSF-MCB) grant MCB-1253444.

