## Supplemental Material for "Self-similar tip growth links exocytosis profile with cell wall shape"

### I. SOLVING THE SECRETION PROFILE $\gamma$ AS AN INITIAL VALUE PROBLEM

Let $\lambda_s^*(s^0) = \lambda_s(s^0) - 1$, $\lambda_\theta^*(s^0) = \lambda_\theta(s^0) - 1$, and $\tilde{\lambda}(s^0) = \lambda_s^*(s^0)/\lambda_\theta^*(s^0)$. From the self-similarity condition Eq.(5) in the main text, and by assuming $\lambda_s^*(s^0) > 0$ and $\lambda_\theta^*(s^0) > 0$ we have

$$\gamma(s^0) = \frac{dr^0/ds^0}{\lambda_\theta^*(s^0)r^0(s^0)} \int_0^{s^0} \lambda_s^*(w)\gamma(w)\, dw. \tag{1}$$

Differentiating through, we obtain

$$\begin{aligned}
\gamma'(s^0) &= \frac{dr^0/ds^0}{\lambda_\theta^*(s^0)r^0(s^0)}\lambda_s^*(s^0)\gamma(s^0) + \Bigg(\frac{\lambda_\theta^*(s^0)r^0(s^0)(d^2r^0/d(s^0)^2)}{(\lambda_\theta^*(s^0)r^0(s^0))^2} \\
&\quad - \frac{(dr^0/ds^0)\left(\lambda_\theta'(s^0)r^0(s^0) + \lambda_\theta^*(s^0)(dr^0/ds^0)\right)}{(\lambda_\theta^*(s^0)r^0(s^0))^2}\Bigg) \int_0^{s^0} \lambda_s^*(w)\gamma(w)\, dw \\
&= \tilde{\lambda}(s^0)\frac{dr^0/ds^0}{r^0(s^0)}\gamma(s^0) + \Bigg(\frac{d^2r^0/d(s^0)^2}{\lambda_\theta^*(s^0)r^0(s^0)} \\
&\quad - \frac{(dr^0/ds^0)(\lambda_\theta'(s^0)r^0(s^0) + \lambda_\theta^*(s^0)(dr^0/ds^0))}{(\lambda_\theta^*(s^0)r^0(s^0))^2}\Bigg)\frac{\gamma(s^0)\lambda_\theta^*(s^0)r^0(s^0)}{dr^0/ds^0} \\
&= \left((\tilde{\lambda}(s^0) - 1)\frac{dr^0/ds^0}{r^0(s^0)} + \frac{d^2r^0/d(s^0)^2}{dr^0/ds^0} - \frac{\lambda_\theta'(s^0)}{\lambda_\theta^*(s^0)}\right)\gamma(s^0).
\end{aligned} \tag{2}$$

Thus, we define

$$f(s^0) = (\tilde{\lambda}(s^0) - 1)\frac{dr^0/ds^0}{r^0(s^0)} + \frac{d^2r^0/d(s^0)^2}{dr^0/ds^0} - \frac{\lambda_\theta'(s^0)}{\lambda_\theta^*(s^0)}, \tag{3}$$

and convert the self-similarity condition to an initial value problem with $\gamma'(s^0) = f(s^0)\gamma(s^0)$ and the initial condition $\gamma(0) = \bar{\gamma}$. Note that $f$ depends only on $\lambda_s$, $\lambda_\theta$, $\lambda_\theta'$, $r^0$, $dr^0/ds^0$, and $d^2r^0/d(s^0)^2$, which means that the secretion is constrained by only the information of the strains up to first-order derivative and the shape up to second-order derivative.

Furthermore, one may verify that

$$\lim_{s^0 \to 0+} f(s^0) = \frac{\lambda_s'(0) - 2\lambda_\theta'(0)}{\lambda_\theta^*(0)} \tag{4}$$

is finite as long as $\lambda_\theta^*(0) = \lambda_s(s^0) - 1 \neq 0$ and the tip is smooth such that $dr^0/ds^0$ is differentiable and $dr^0/ds^0|_{s^0=0} = 1$. We define $f(0) = \lim_{s^0\to 0+} f(s^0)$. Then there exists a unique solution $\gamma(s^0) = \overline{\gamma}\exp(\int_0^{s^0} f d\omega)$ on some interval $[0, \Sigma)$ as long as $r^0 \neq 0, dr^0/ds^0 \neq 0, \lambda_s \neq 1$ such that $f$ is continuous on $[0, \Sigma)$.

The above analysis gives rise to criteria for detecting whether the distribution of secretion is monotonic (Theorem 1) and determining if there exists a local maximum of $\gamma$ at the tip (Theorem 2).

**Theorem 1.** *Given $\lambda_s$ continuous and $\lambda_\theta$, $r^0$ and $dr^0/ds^0$ continuously differentiable, the secretion distribution $\gamma$ from Eq.(1) and the condition $\gamma(0) = \overline{\gamma} > 0$ is strictly monotonic on the interval $[0, \Sigma] \subset D = \{0\} \cup \{s^0 | r^0 \neq 0, dr^0/ds^0 \neq 0, \lambda_s \neq 1\}$ iff $f(s^0)$ from Eq.(3) never vanishes away from the tip.*

*Proof.* The forward direction is obvious using proof by contradiction. Now we prove the backward direction. Given the uniqueness property, the solution to $\gamma'(s^0) = f(s^0)\gamma(s^0)$ and $\gamma(0) = \overline{\gamma}$ takes the form $\gamma(s^0) = \overline{\gamma}\exp(\int_0^{s^0} f d\omega)$. If $f$ never vanishes, we have either $f > 0$ or $f < 0$. Thus, $\gamma(s^0)$ either monotonically increases (given $f > 0$) or monotonically decreases (given $f < 0$).

□

**Theorem 2.** *Given $\lambda_\theta$, $\lambda'_\theta$, $\lambda'_s$, $\gamma$ continuous and $\lambda_\theta > 1$ and $\gamma(0) > 0$, the secretion distribution $\gamma$ has a local maximum at the tip $s^0 = 0$ if*

$$\frac{\lambda'_s(0) - 2\lambda'_\theta(0)}{\lambda_\theta(0) - 1} < 0 \tag{5}$$

*and has a local minimum at the tip point if*

$$\frac{\lambda'_s(0) - 2\lambda'_\theta(0)}{\lambda_\theta(0) - 1} > 0 \tag{6}$$

*Proof.* We will only prove the first statement. By continuity, there exists $\delta > 0$ such that $f(s^0) < 0$ for $0 \leq s^0 < \delta$. By integrating $\gamma' = f\gamma$ , we obtain

$$\gamma(s^0) - \gamma(0) = \int_0^{s^0} f\gamma d\omega < 0 \tag{7}$$

for any $0 < s^0 < \delta$. So $\gamma$ has one local maximum at $s^0 = 0$. □

### II. THE CUBIC-SPLINE SOLUTIONS

The simulation for elastic deformation is based on the cubic-spline solutions of $r(s)$, $z(s)$, and $s(s^0)$ given the cubic splines of coordinates $r^0(s^0)$ and $z^0(s^0)$. The steps of this procedure are summarized as follows:

1. Parametrize the initial, unturgid outline by $(z^0(s^0), r^0(s^0))$ and discretize this outline by $N$ marker points $(z_i^0, r_i^0)$. Interpolate between the marker points $(z_i^0, r_i^0)$, using cubic splines.

2. Finding the initial guess of the steady-state turgid configuration $(z_i, r_i)$: approximate the steady-state turgid configuration $(z_i, r_i)$ by displacing the marker points $(z_i^0, r_i^0)$ according to $d\vec{R}/dt = \vec{F}$ where $\vec{R}$ is the vector concatenated by the $z$- and $r$-coordinates of all marker points. The components of the force vector $\vec{F}$ along $z$- and $r$-direction on each marker point are computed from Eq.(6) in the main text. In Eq.(6) in the main text, the stresses $\sigma_s^i$ and $\sigma_\theta^i$ are computed through Eqs.(7),(8) and (3) in the main text where the elastic stretches $\lambda_s^i$ and $\lambda_\theta^i$ are computed based on the cubic splines of $s(s^0)$, $r(s)$, and $r^0(s^0)$. The components of $\vec{F}$ related to the boundary marker points ($i = 1$ and $i = N$) rely on Eqs.(9) and (10) in the main text respectively. The points $(z_i, r_i)$ are updated until the force residue is below a certain threshold. Notice that for each iteration, the cubic-spline coefficients of $s(s^0)$ and $r(s)$ are updated while $r^0(s^0)$ maintains the same.

3. Finding the solution of the steady-state turgid configuration $(z_i, r_i)$: based on the initial guess of $(z_i, r_i)$, use Newton's method to solve iteratively for the configuration of the points $(z_i, r_i)$ which leaves machine-zero force residue. Again, for each iteration, the cubic-spline coefficients are updated. Given both unturgid and turgid coordinates, we can solve elastic stretches $\lambda_s, \lambda_\theta$ by Eq.(3) in the main text.

For the inference of the exocytosis $\gamma$ over the cell outline, we use directly Eq.(5) in the main text which only requires numerical integration.

For the simulation of cell wall extension in Fig.2c in the main text, we update the unturgid wall data $s_i^0$, $r_i^0$ for each marker point using the directional expansion distributions according to Eqs. (2) and (4) from the main text by a linear approximation $r_{new}^0 = r_{old}^0 + \Delta t \gamma_\theta r_{old}^0$, and similarly for $s^0$, where $\Delta t$ is a small time step. We repeat step 1-3 at every update of the unturgid wall configuration $s_i^0$, $r_i^0$.

### III. FITTING THE DATA INTO A CANONICAL SHAPE

#### A. Data preprocessing

The cell shape data were provided to us as a set of outlines, each given by a sequence of $X$ and $Y$ coordinates of points on the cell outline. We first truncated each sequence of $X$ and $Y$ coordinates to remove any region beyond the point where the shank width stops increasing. Then we interpolated $X$ and $Y$ with cubic splines and finally placed marker points on the interpolated outline uniformly with respect to arclength, making an arbitrarily fine discretization. The resolution of the discretization has little effect on the inferred secretion (not shown).

#### B. Cell shape fitting

Our initial attempt at characterizing the inferred secretion under a wide variety of tip geometries used a one-parameter family of shapes comprising a cylindrical pipe with ellipsoidal endcap [1]. However, this family of shapes does not fit the available data (moss *Physcomitrium patens* protonemata and *Medicago truncatula* root hairs) well. See Fig. S 1 for example.

Then, we adopt a one-parameter family of cell outline shapes given by so-called hyphoid shapes [2]. These shapes are defined by the equation

$$z = \frac{\pi r}{a}\cot(\pi r). \tag{8}$$

Here $a$ is a parameter controlling the shape of the tip; larger values of $a$ correspond to a flatter tip, and in the limit as $a \to \infty$, the hyphoid shape approaches a rectangle.

The hyphoid function fits the data accurately in a least-squares sense with a horizontal translation $z = \frac{\pi r}{a}\cot(\pi r) - \frac{1}{a}$ (such that $z(r) = 0$). See Fig. S 1 for example. We observed that fitted moss protonema and root hair outlines qualitatively recover the curvature distributions of moss protonemata in [3] and root hairs in [4] (not shown).

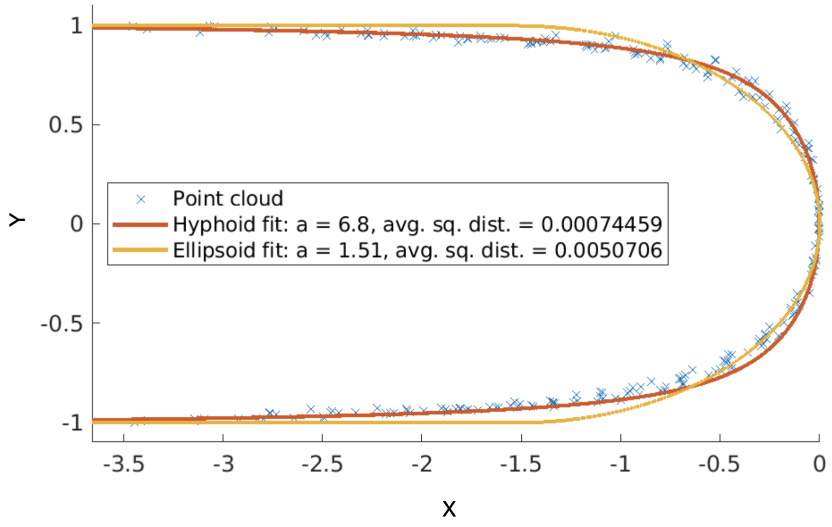


FIG. S1: Fitting the point cloud of root hair shapes with hyphoid and ellipsoid shapes. The fitting is done by minimizing the sum of squared distances between the points and the fit curve.

To set up the unturgid shape for our simulations, we truncate the cell outline at $z = -5$ from Eq.(8) to ensure that we capture enough of the shape to reliably observe the tip behavior. Although the choice to truncate at $z = -5$ is arbitrary, its effect on the inferred secretion is negligible (not shown).

### IV. INSENSITIVITY TO ELASTIC LAWS

To demonstrate that the non-monotonicity of the exocytosis profile is not a result of the assumption of elastic properties, Figs. S 2a and S2b show the inferred secretion for the unturgid shapes in Fig. 2 of the main text under the assumptions of softer nonlinear elastic material and linear elastic material.

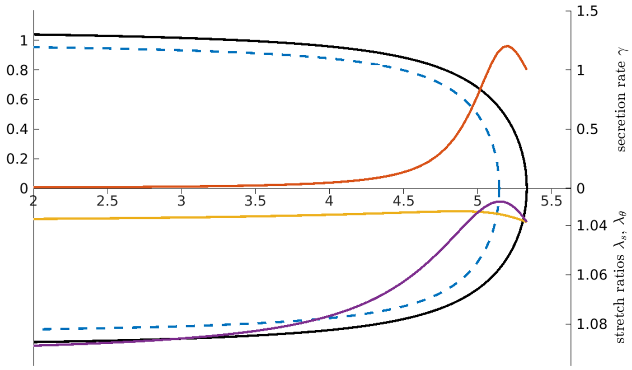


(a) Deformation and inferred secretion for the same unturgid shape as Fig. 2 in the main text, with softer material ($K_h = \mu_h = 6$).

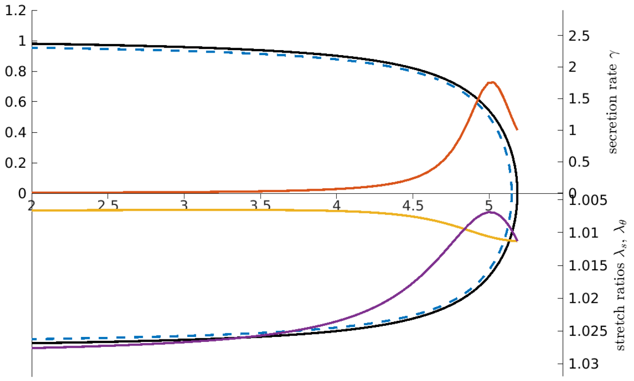


(b) Deformation and inferred secretion for the same unturgid shape as Fig. 2 in the main text, under linear elasticity ($\nu = 0.3$, $E_h = 30$).

FIG. S2: The secretion distribution $\gamma$ is qualitatively preserved when the mechanical parameters (a) or the elastic law is changed (b).

### V. DETAILS OF FIG. 3 IN THE MAIN TEXT

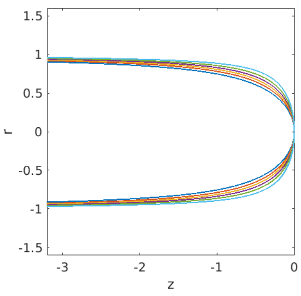


(a) Unturgid shapes

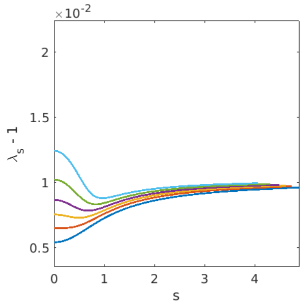


(b) Overlaid plots of $\lambda_s$

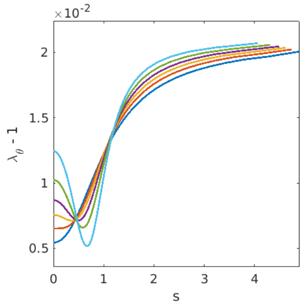


(c) Overlaid plots of $\lambda_\theta$

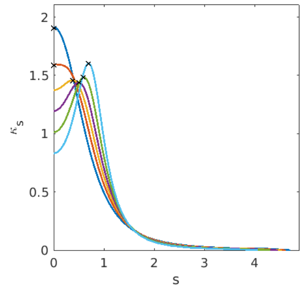


(d) Overlaid plots of $\kappa_s$

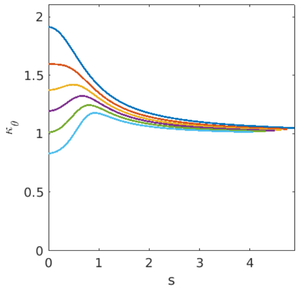


(e) Overlaid plots of $\kappa_\theta$

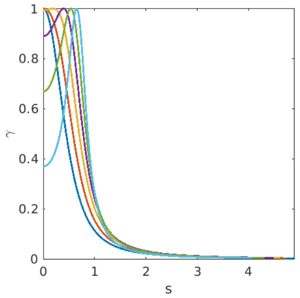


(f) Overlaid plots of $\gamma$

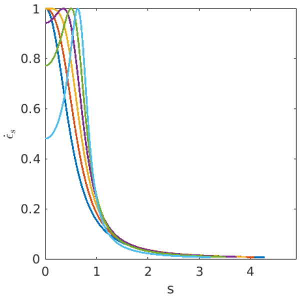


(g) Overlaid plots of $\dot{\epsilon}_s$

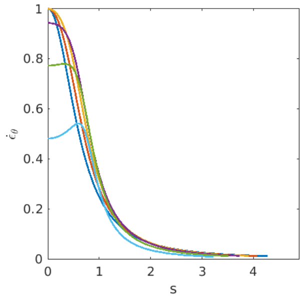


(h) Overlaid plots of $\dot{\epsilon}_\theta$

FIG. S3: This range of tip shapes captures a large portion of the parameter space $R$ shown in Fig. 3 of the main text. The selected hyphoids correspond to $R$ approximately between 0.55 and 1.2. In subfigure (d), the location of maximum meridional curvature for each turgid shape is marked with a black $\times$.The maximal curvature location transition is continuous as the hyphoid shape transitions from being tapered to flattened at the tip. The transition of ellipse shapes cannot achieve this continuous transition.

See Fig. S 3 for the change of profiles of secretion $\gamma$, elastic stretches $\lambda_s$ and $\lambda_\theta$, curvatures $\kappa_s$ and $\kappa_\theta$, and wall extension rates $\dot{\epsilon}_s$ and $\dot{\epsilon}_\theta$ as the cell shape transitions from being tapered to flat at the tip end. We normalize the rates

($\gamma$, $\dot{\epsilon}_s$, and $\dot{\epsilon}_\theta$) by the rate maximum and normalize curvatures ($\kappa_s$ and $\kappa_\theta$) by $1/r(s_b)$, the inverse of the maximum radial width of the cells. In each of our simulations, $r(s_b)$ is approximately equal to 1.


[1] P. Fayant, O. Girlanda, Y. Chebli, C.-É. Aubin, I. Villemure, and A. Geitmann, Finite element model of polar growth in pollen tubes, The Plant Cell **22**, 2579 (2010).

[2] S. Bartnicki-Garcia, F. Hergert, and G. Gierz, Computer simulation of fungal morphogenesis and the mathematical basis for hyphal (tip) growth, Protoplasma **153**, 46 (1989).

[3] D. Chelladurai, G. Galotto, J. Petitto, L. Vidali, and M. Wu, engInferring lateral tension distribution in wall structures of single cells, European physical journal plus **135** (2020).

[4] J. Dumais, S. R. Long, and S. L. Shaw, The Mechanics of Surface Expansion Anisotropy in Medicago truncatula Root Hairs, Plant Physiology **136**, 3266 (2004), https://academic.oup.com/plphys/article-pdf/136/2/3266/36005831/plphys_v136_2_3266.pdf.
